# BTLA and PD-1 employ distinct phosphatases to differentially repress T cell signaling

**DOI:** 10.1101/669812

**Authors:** Xiaozheng Xu, Amitkumar Fulzele, Yunlong Zhao, Zijun Wu, Yanyan Hu, Yong Jiang, Yanzhe Ma, Haopeng Wang, Guo Fu, Eric Bennett, Enfu Hui

## Abstract

T cell-mediated destruction of tumors and virus-infected cells is restricted by co-inhibitory receptors such as programmed cell death protein 1 (PD-1). Monoclonal antibodies blocking PD-1 have produced impressive clinical activity against human cancers, but durable response is limited to a minority of patients. Previous results suggest that B and T lymphocyte attenuator (BTLA), a co-inhibitory receptor structurally related to PD-1, may contribute to the resistance to PD-1 targeted therapy and co-blockade of BTLA can enhance the efficacy of anti-PD-1 immunotherapy. However, the biochemical mechanism by which BTLA represses T cell activity and to what extent the mechanism differs from that of PD-1 is unknown. Here we examine differences in the ability of BTLA and PD-1 to recruit effector molecules and regulate T cell signaling. We show that PD-1 and BTLA recruit different tyrosine phosphatases to regulate either CD28 or T cell antigen receptor (TCR)-signaling cascades. Our data reveal unexpected disparities between two structurally related immune checkpoints and two phosphatase paralogs.

## INTRODUCTION

T cell activation is governed by both antigen-specific signals from TCR and antigen-nonspecific signals through coreceptors. The relative strength of these signaling pathways – with some positively regulate T cell activation (co-stimulatory), and others repressing the T cell activation (co-inhibitory) - are critical in shaping the overall immune response (Chen and Flies, 2013). Several co-signaling receptors belong to the B7 family of the immunoglobulin (Ig) superfamily. Among these, CD28 is a central co-stimulatory receptor that upon binding to its ligands B7-1 or B7-2 (Lenschow et al., 1996), delivers essential positive signals for full activation of naïve T cells (Lanzavecchia et al., 1999). PD-1 and BTLA are both co-inhibitory receptors that attenuate T cell activation (Carreno and Collins, 2003; Freeman et al., 2000; Nishimura and Honjo, 2001; Watanabe et al., 2003).

PD-1 is absent on naïve T cells, and transiently induced upon TCR activation to restrain excessive T-cell-mediated tissue damage (Keir et al., 2008). In contrast, BTLA is abundant on naïve T cells but downregulated during T cell development and differentiation (Baitsch et al., 2012; Derre et al., 2010; Watanabe et al., 2003). BTLA downregulation appears to be essential to the function of effector T cells (Derre et al., 2010; Pasero and Olive, 2013). PD-1 has two known ligands in the B7 family (Greenwald et al., 2005; Sharpe and Freeman, 2002): the lower affinity yet broadly expressed programmed death-ligand 1 (PD-L1) (Freeman et al., 2000; Taube et al., 2012), and the higher affinity, more restrictly expressed programmed death-ligand 1 (PD-L2) (Cheng et al., 2013). Notably, the best studied ligand for BTLA, herpes virus entry mediator (HVEM) (Compaan et al., 2005; Gonzalez et al., 2005; Sedy et al., 2005), is a member of the TNF receptor (TNFR) family rather than B7 family (Croft, 2003; Morel et al., 2000).

Normally acting as “checkpoints” or “molecular brakes” to prevent overreactive T cells (Marraco et al., 2015), co-inhibitory receptors can be hijacked by tumors and viruses to escape immune destruction (Baitsch et al., 2011; Mellman et al., 2011; Pardoll, 2012; Pauken and Wherry, 2015; Sharma and Allison, 2015). Elevated expression of PD-1, BTLA and several other co-inhibitory molecules in cancer patients or mouse tumor models often signify the hypofunction, or “exhaustion” of T cells (Jiang et al., 2015; Thommen and Schumacher, 2018). Antibodies that block the PD-1 pathway have proven impressive clinical activities against several human cancers in a fraction of patients (Hamid et al., 2013; Herbst et al., 2014; Powles et al., 2014; Rizvi et al., 2015; Topalian et al., 2012). Evidence suggest that compensatory upregulation of other co-inhibitory receptors in response to PD-1 blockade therapy might contribute to the observed resistance to PD-1 inhibitors (Stecher et al., 2017). Combination immunotherapy that blocks PD-1 and cytotoxic T lymphocyte antigen-4 (CTLA-4) has demonstrated superior clinical activity in melanoma patients (Larkin et al., 2015; Wolchok et al., 2013). In addition, combination blockade of PD-1 and BTLA leads to superior T cell proliferation and cytokine production than mono-blockade of PD-1 (Stecher et al., 2017). Further, BTLA/PD-1 co-expression is required for the exhaustion of human hepatocellular carcinoma infiltrated CD4^+^ T cells (Zhao et al., 2016), consistent with an important role of BTLA in repressing the anti-tumor activity of T cells.

Both PD-1 and BTLA contain an Ig-like ectodomain, a single transmembrane segment, and an intracellular tail. The tail of PD-1 contains two tyrosines, Y223 and Y248, embedded in an immunoreceptor-tyrosine-inhibitory motif (ITIM) and an immunoreceptor-tyrosine-switch-motif (ITSM), respectively. Analogous to PD-1, the cytoplasmic tail of BTLA contain both an ITIM (surrounding Y257) and an ITSM (surrounding Y282). There are also two additional tyrosines (Y226 and Y243) N terminal to the ITIM (Chemnitz et al., 2006). Phospho-peptide pull down assays suggest that two additional tyrosine residues (Y226 and Y243) N-terminal to ITIM might bind GRB2 (Gavrieli and Murphy, 2006), an adaptor protein that might nucleate additional signaling events.

Engagement of PD-1 with either PD-L1 or PD-L2 triggers phosphorylation of both Y223 and Y248, and recruitment of the protein tyrosine phosphatase Src-homology-2 (SH2) domain-containing phosphatase 2 (SHP2), and possibly SH2 domain-containing phosphatase 1 (SHP1), through their tandem SH2 (tSH2) domains (Chemnitz et al., 2004). PD-1 associated SHP2 was shown to dephosphorylate co-stimulatory receptors CD28 and CD226, with a much weaker effect on the phosphorylation of the immunoreceptor-tyrosine-based-activation motif (ITAM) of TCR (Hui et al., 2017; Wang et al., 2018). However, a recent study showed that SHP2 might be dispensable for PD-1 to contribute to T cell exhaustion (Rota et al., 2018), raising the question of whether SHP2 is the major effector of PD-1. It is also unclear whether and how SHP1 contributes to PD-1 function.

Much less is known about the mechanism of BTLA signaling and how BTLA signaling differs from PD-1 signaling. Co-immunoprecipitation experiments in transfected cell lines suggest that HVEM:BTLA interaction triggers BTLA phosphorylation and recruitment of SHP1 and SHP2 (Gavrieli et al., 2003; Sedy et al., 2005). However, these findings contradict to other studies showing that BTLA recruits SHP1, but not SHP2 in HVEM-stimulated primary CD4+ T cells (Chemnitz et al., 2006). Paradoxically, this study also indicates that mutations that abrogate SHP1 recruitment do not affect the ability of BTLA to suppress interleukin-2 (IL-2) secretion. Hence, further studies are needed to define the BTLA signaling pathway.

In this study, we have systematically compared BTLA and PD-1 signaling in an antigen-presenting cell (APC)-T cell co-culture system. We find that upon ligation, PD-1 recruits SHP2 but not SHP1 whereas BTLA preferentially recruits SHP1. Moreover, the PD-1:SHP2 complex dephosphorylates the co-stimulatory receptor CD28 but not TCR under the conditions tested. Similarly, the BTLA:SHP2 complex dephosphorylates CD28 but not TCR. This is in contrast to BTLA:SHP1 complex which potently dephosphorylates both TCR and CD28. These results reveal an unexpected functional disparity between two structurally related immune checkpoint receptors, as well as between two phosphatase paralogs that are implicated in many signaling pathways.

## RESULTS AND DISCUSSION

### Both BTLA and PD-1 inhibit IL-2 production from T cells

We first sought to establish a cell culture system to compare PD-1 and BTLA signaling in parallel. To this end, we used Jurkat T cells as the responder cells and Raji B cells as the APCs. Raji B cells preloaded with the superantigen staphylococcal enterotoxin E (SEE) stimulate TCR signaling in Jurkat T cells, upon their physical contact, due to the ability of SEE to crosslink MHC and TCR (Choi et al., 1989). Meanwhile, the Raji-Jurkat cell interaction also triggers CD28 co-stimulatory signaling due to the expression of B7 ligand molecules on Raji cells (Tatsumi et al., 1997). Raji and Jurkat cells are both excellent hosts for lentiviral transduction, allowing stable expression of ligands or receptors of interest.

Using this co-culture system, we recently demonstrated that CD28 is a primary target of PD-L1:PD-1 mediated inhibition (Hui et al., 2017). However, we did not address if PD-1 signals exclusively through the two tyrosine motifs (i.e., ITIM and ITSM) or additional mechanisms are involved. To this end, we established two Jurkat lines that express similar levels of either PD-1 wild-type (WT) or mutant PD-1 in which both intracellular tyrosines were mutated to phenylalanine [denoted as PD-1 (FF)] (**Fig 1A** and **B**). Both versions of PD-1 were fused with a C-terminal mGFP tag to facilitate immunoprecipitation experiments described below, and are referred as PD-1 (WT)-mGFP and PD-1 (FF)-mGFP, respectively. As expected, upon co-culturing with SEE-loaded PD-L1-transduced Raji cells, Jurkat [PD-1 (WT)-mGFP] secreted much less IL-2 than the mock-transduced Jurkat (WT) (**Fig 1C**). Replacement of PD-1 (WT)-mGFP with equal levels of PD-1 (FF)-mGFP completely restored the IL-2 secretion, indicating that PD-1 primarily, if not exclusively, signals through its two intracellular tyrosine motifs.

**Figure 1.**
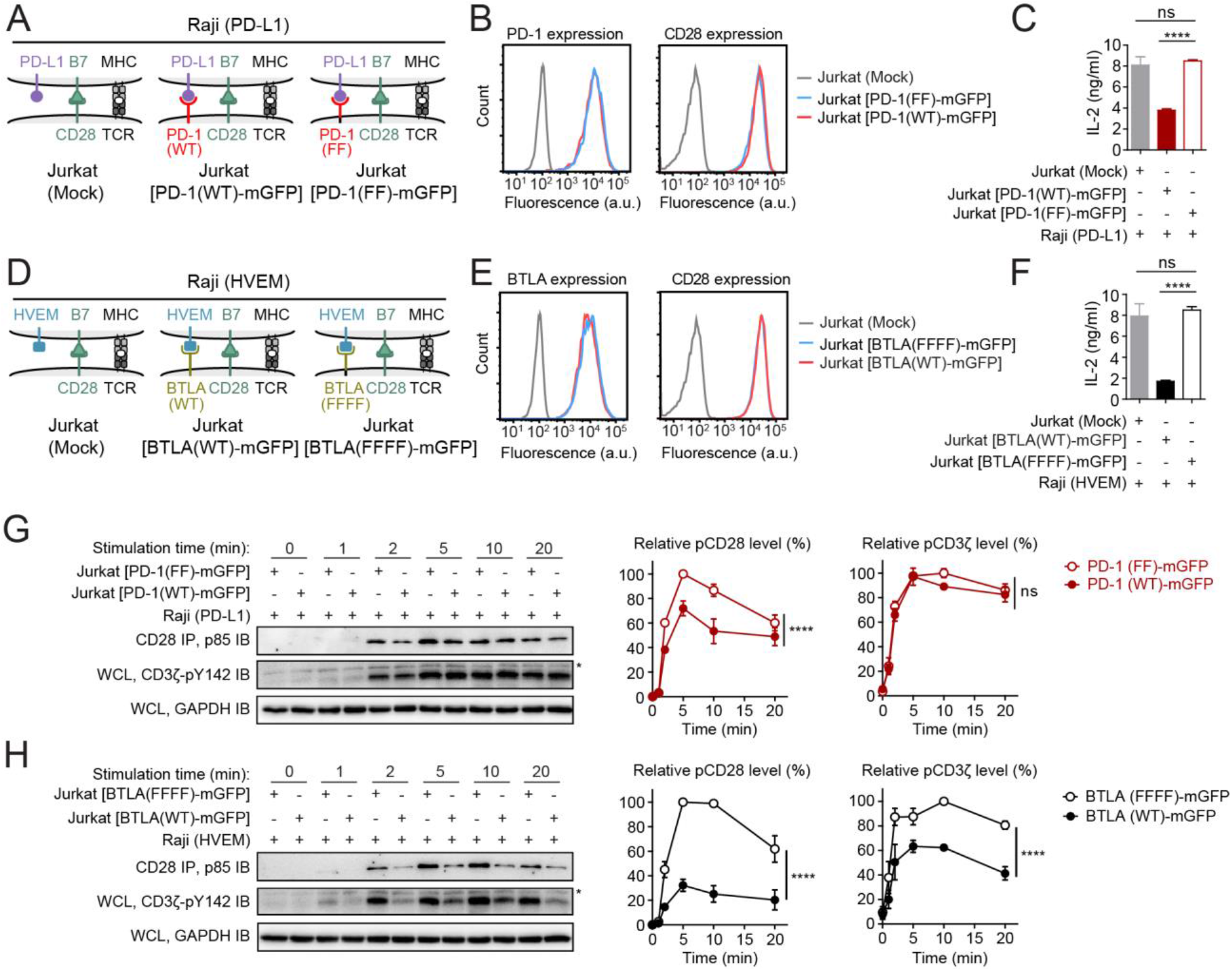
PD-1 inhibits CD28 phosphorylation while BTLA inhibits both TCR and CD28 phosphorylation. **(A)** Cartoon illustrating an intact cell assay in which mock transduced, PD-1 (WT)-mGFP or PD-1 (FF)-mGFP transduced Jurkat T cells were stimulated with PD-L1 transduced Raji B cells preloaded with SEE antigen. **(B)** Flow cytometry histograms showing PD-1 and CD28 surface expressions in the indicated Jurkat T cells. a. u. denotes arbitrary units. **(C)** Bar graphs summarizing IL-2 levels in the medium of the indicated Jurkat-Raji co-cultures (see Methods). (**D-F**) Same as A-C except PD-1 (WT)-mGFP, PD-1 (FF)-mGFP, and PD-L1 were replaced with BTLA (WT)-mGFP, BTLA (FFFF) -mGFP and HVEM respectively. **(G, H)** Shown on the left are representative immunoblots showing the phosphorylation of CD3ζ (anti-pY142) and CD28 (co-IP’ed p85) in the indicated co-cultures, with the cells lysed at the indicated times after the initial contact (see Methods). GAPDH IB indicates the input of each sample. Non-specific bands were labeled by asterisks. WCL, whole cell lysate. Shown on the right are quantification graphs of CD28 and CD3ζ phosphorylations. Data were normalized to the highest phosphorylation level under PD-1 FF or BTLA FFFF condition, respectively. Data in this figure are presented as means ± SD from three independent measurements. ****P < 0.0001; ns, not significant; Two-way ANOVA (analysis of variance).

Likewise, to study HVEM:BTLA signaling, we transduced Jurkat cells, which do not express BTLA endogenously (**Fig S1A**), with either BTLA (WT)-mGFP or a signaling-deficient BTLA in which all four intracellular tyrosines (Y226, Y243, Y257, Y282) were mutated to phenylalanine (BTLA (FFFF)-mGFP) (**Fig 1D**). According to flow cytometry, the two versions of BTLA were expressed at similar levels and neither affected the expression of CD28 (**Fig 1E**). Upon co-culturing with SEE-loaded HVEM-transduced Raji cells, Jurkat cell expressing BTLA (WT)-mGFP produced significantly lower IL-2 than Jurkat cells expressing BTLA (FFFF)-mGFP. Importantly, Jurkat cells expressing the mutant BTLA produced indistinguishable levels of IL-2 compared to mock-transduced BTLA-negative Jurkat cells (**Fig 1F**). This result indicates that the four intracellular tyrosines of BTLA are necessary for BTLA to inhibit IL-2 production of T cells.

### BTLA inhibits both TCR and CD28 while PD-1 inhibits CD28 phosphorylation

Having established the PD-1 and BTLA effects on IL-2 secretion, we next compared their signaling at the receptor level. We recently reported that PD-1 preferentially inhibits the phosphorylation of CD28 over CD3ζ subunit of the TCR complex (Hui et al., 2017). Consistent with this finding, a direct comparison of Jurkat cells expressing PD-1 (WT)-mGFP and Jurkat cells expressing PD-1 (FF)-mGFP confirmed our previous finding that PD-1 inhibits the phosphorylation of CD28 but not CD3ζ (**Fig 1G**).

We next examine if BTLA signaling differs from PD-1 signaling with regards to CD3ζ and CD28 phosphorylation. We conjugated either Jurkat cells expressing BTLA (WT)-mGFP or BTLA (FF)-mGFP with SEE-loaded HVEM+ Raji cells and collected cells over time. Subsequent immunoblotting (IB) of co-culture lysates revealed that conjugation with Raji cells induced a time-dependent phosphorylation of both CD3ζ and CD28, with the latter reported by the co-immunoprecipitated (IP) p85, which binds CD28 only if CD28 is phosphorylated (see **Methods**). Notably, phosphorylation of both CD3ζ and CD28 in Jurkat cells expressing BTLA (WT)-mGFP were substantially weaker compared to cells expressing mutant BTLA (**Fig 1H**) over the entire time course. These results demonstrate that unlike PD-1, BTLA potently inhibits both TCR and CD28 phosphorylation.

BTLA expression in Jurkat cells, compared to PD-1 expression, results in a more robust inhibition of IL-2 production and CD28 phosphorylation than does PD-1 (**Fig 1F** versus **1C, 1H** versus **1G**). This observation could, in principle, be due to a higher BTLA expression than PD-1 expression. To clarify this issue, we quantified BTLA (WT)-mGFP and PD-1 (WT)-mGFP expressions using purified GFP as a standard (**Fig S2**). This experiment revealed that PD-1 expression was 3.4-fold higher than BTLA in the transduced Jurkat cells. Furthermore, we also examined the expression of their ligands in the respective Raji APCs, and found that PD-L1 was more highly expressed than HVEM (**Fig S1B**). The lower expression of BTLA/HVEM than PD-1/PD-L1, in conjunction with the stronger inhibitory effects of BTLA presented in **Fig 1**, suggests that BTLA is an intrinsically more potent inhibitory receptor than PD-1. This finding is surprising in the context of more severe autoimmunity associated with PD-1 knockout (Iwai et al., 2017; Nishimura et al., 1999; Nishimura et al., 2001) and the better clinical activities of PD-1 inhibitors (Hamid et al., 2013; Herbst et al., 2014; Powles et al., 2014; Rizvi et al., 2015; Topalian et al., 2012). It is likely that the spatiotemporal expression of these two receptors and their ligands influence their functional outcomes.

### BTLA recruits both SHP1 and SHP2 while PD-1 recruits only SHP2

To investigate the molecule mechanism underlying the different potency and specificities of BTLA and PD-1-mediated suppression of T cell activation, we examined the landscape of intracellular effector proteins recruited to either BTLA or PD-1. We used GST-tagged, prephosphorylated tail of either PD-1 or BTLA to pull down proteins from Jurkat T cell lysates to define their interactomes using mass spectrometry (Peled et al., 2018). One-side volcano plots revealed that PD-1 and BTLA both pulled down SHP1 and SHP2. Several other SH2 proteins are also among the top hits. Interestingly, ZAP70 co-precipitated with BTLA but not PD-1, whereas SAP and p85α, the regulatory subunit of PI3 kinases, co-precipitated with PD-1 (**Fig 2A**). However, GRB2, which was suggested to bind BTLA (Gavrieli and Murphy, 2006; Ritthipichai et al., 2017), was not detected in this assay.

**Figure 2.**
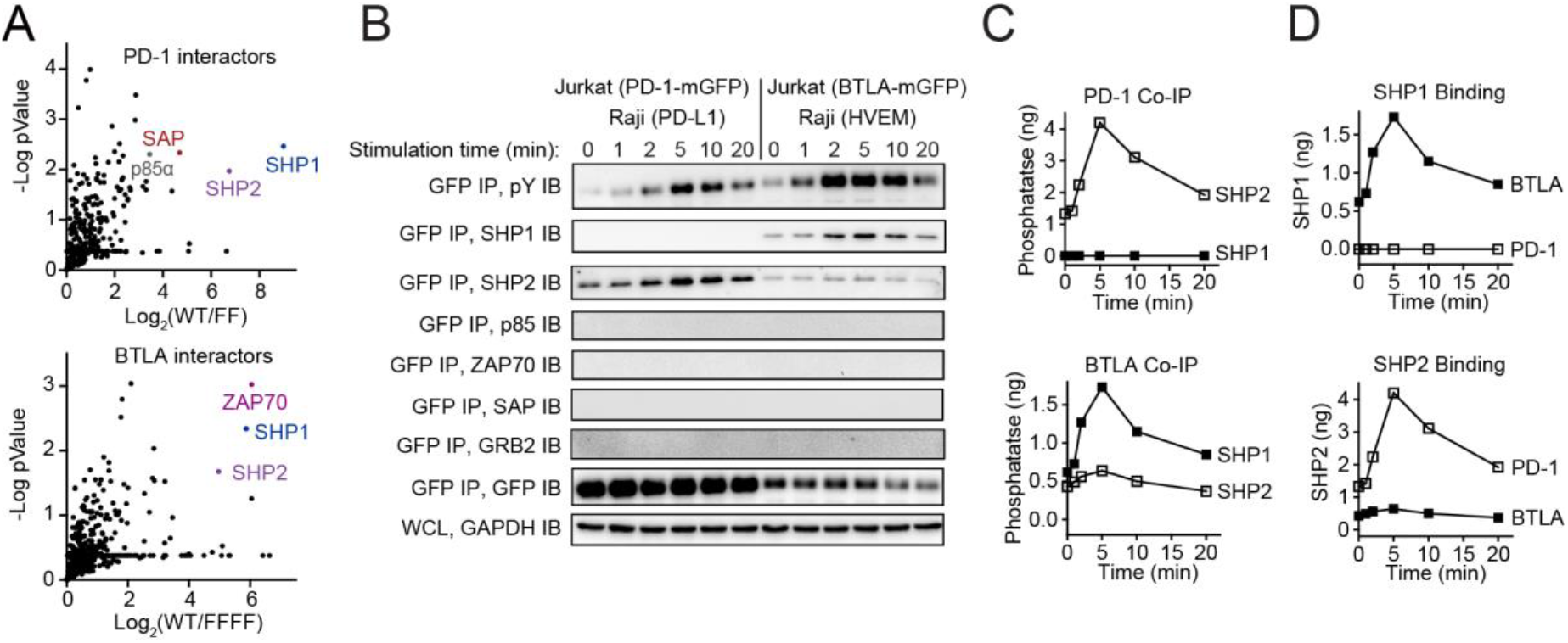
PD-1 recruits only SHP2 while BTLA recruits both SHP1 and SHP2. **(A)** One-side volcano plot showing proteins that potentially interact with the intracellular tail of PD-1 or BTLA. SH2 domain-containing candidates are highlighted in colors. GST-tagged, prephosphorylated PD-1 or BTLA tail was used as the bait to pull down proteins from Raji-stimulated Jurkat lysates, and proteins identities determined by Mass Spec approach (see Methods). **(B)** Representative immunoblots showing the levels of tyrosine phosphorylation (pY IB) of immunoprecipitated PD-1-mGFP or BTLA-mGFP (GFP IP), as well as the co-precipitated SHP1, SHP2, p85, ZAP70, SAP and GRB2 from the indicated co-culture lysates, with the time at which the cell lysis occurred indicated (see Methods). GAPDH IB indicates the input of each sample. **(C)** Co-precipitated SHP1 and SHP2 in (B) were quantified using purified recombinant SHP1/SHP2 standards and plotted as SHP1 versus SHP2 recruitments to PD-1 or BTLA, or **(D)** PD-1 versus BTLA in recruiting either SHP1 or SHP2.

To validate these interactors in a more physiological setting, we co-IP’ed either PD-1-GFP or BTLA-GFP from Raji-Jurkat co-cultures, lysed at the indicated time points after conjugation, and detected the potential interactors using immunoblots (IB) (**Fig 2B**). These experiments showed that PD-1 selectively recruits SHP2 but not SHP1, while BTLA strongly prefers SHP1 over SHP2 (**Fig 2B** and **C**). Moreover, phosphatase recruitment to both PD-1 and BTLA occurred in a time-dependent manner, with recruitment peaking at 5 minutes, and decreasing to basal levels by 20 minutes after Raji-Jurkat conjugation. From 2 to 10 minutes, BTLA recruited 2-3 folds more SHP1 than SHP2. By contrast, ZAP70, SAP, p85 and GRB2 were undetectable in either PD-1 or BTLA precipitates over the entire time course (**Fig 2B**), indicating that even some of these proteins may interact with PD-1 or BTLA under certain circumstances, the affinities are likely very weak. Because GST exists predominantly as a dimer (Fabrini et al., 2009), we speculate that the GST-mediated dimerization of the bait proteins might have facilitated these otherwise weak, and perhaps non-physiological interactions.

We then re-plotted the data in **Fig 2C** to directly compare the abilities of PD-1 and BTLA to recruit either SHP1 or SHP2 (**Fig 2D**). Clearly, SHP1 was recruited by BTLA but not by PD-1. By contrast, SHP2 associated with both BTLA and PD-1, but with a ~10-fold preference to PD-1. It is possible, but unlikely, that the 3.4-fold higher expression of PD-1 compared to BTLA in the transduced Jurkat cells (**Fig S2**) would account for the 10-fold difference in SHP2 recruitment. Rather, these results suggest that SHP2 has an intrinsic binding preference for PD-1 over BTLA.

### The intracellular tails of BTLA and PD-1 dictate their specificities

Due to the structural similarities of the intracellular tails of PD-1 and BTLA, their ability to differentially recruit SHP1 and SHP2 was surprising. In principle, these distinctions could be due to either intracellular tails or ectodomains which engage different ligands, i.e., PD-L1 and HVEM respectively. To dissociate the intracellular and extracellular contributions, we created Jurkat cells expressing a PD-1:BTLA chimera, which fuses PD-1 ecto and transmembrane domains with BTLA cytoplasmic tail and a C-terminal mGFP tag (PD-1^ECTO-TMD^-BTLA^CYTO^-mGFP) (**Fig 3A**). Notably, the expression level of this chimera receptor in its host Jurkat cells is similar to PD-1 (WT)-mGFP and PD-1 (FF)-mGFP in their respective host Jurkat cells (**Fig 3B**). Moreover, this chimera receptor allowed us to trigger BTLA intracellular signaling via PD-L1, and therefore directly compare the contribution of the PD-1 and BTLA intracellular tails in the Raji-Jurkat co-culture assay. Upon conjugating the respective Jurkat cells with SEE-loaded PD-L1+ Raji cells, both PD-1 (WT)-mGFP and PD-1^ECTO-TMD^-BTLA^CYTO^-mGFP were tyrosine phosphorylated (**Fig 3C**, GFP IP, pY IB) and recruited SHP2 (**Fig 3C**, GFP IP, SHP2 IB). By contrast, SHP1 was recruited by the chimera PD-1^ECTO-TMD^-BTLA^CYTO^-mGFP but not by PD-1 (WT)-mGFP (**Fig 3C**, GFP IP, SHP1 IB). This result demonstrates that the tail of BTLA is sufficient to recruit SHP1, even if it is fused to the PD-1 ectodomain.

**Figure 3.**
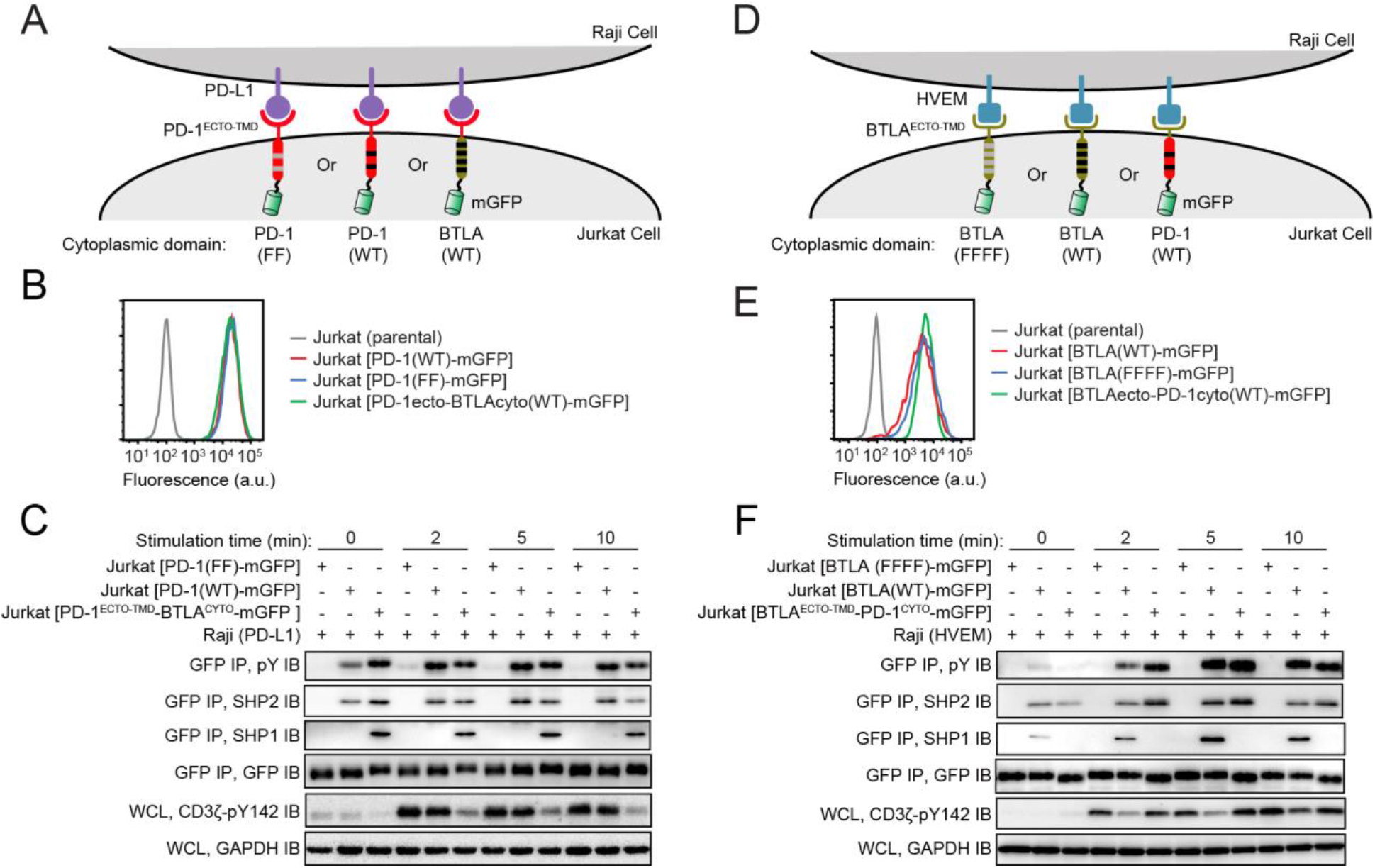
The biochemical specificities of PD-1:BTLA chimeras is dictated by their tails. **(A)** Cartoon illustrating a Raji-Jurkat co-culture assay in which PD-1 (FF)-mGFP, PD-1 (WT)-mGFP, or PD-1ecto-BTLAcyto (WT)-mGFP chimera–transduced Jurkat T cells were stimulated with PD-L1 transduced Raji B cells preloaded with SEE antigen. **(B)** Flow cytometry histograms showing PD-1 antibody staining of indicated Jurkat T cells. a. u. denotes arbitrary units. **(C)** Representative immunoblots of tyrosine phosphorylation (pY IB) and the associated SHP1/SHP2 of the IP’ed GFP-tagged receptors. Immunoblot of CD3ζ (anti-pY142) for the WCL is also shown to reflect how the indicated GFP-tagged receptors affect TCR phosphorylation (see **Methods**). GFP was immunoblotted to show the input of mGFP-tagged proteins. GAPDH was immunoblotted to show the input of cell lysates. (**D-F**) Same as A-C except replacing PD-1 (FF)-mGFP, PD-1 (WT)-mGFP and PD-1ecto-BTLAcyto (WT)-mGFP chimera with BTLA (FFFF)-mGFP, BTLA (WT)-mGFP and BTLAecto-PD-1cyto (WT)-mGFP chimera, respectively, on the Jurkat side. Accordingly, PD-L1 was replaced with HVEM on the Raji side.

In the reciprocal experiment, we generated Jurkat cells expressing the BTLA:PD-1 chimera, in which BTLA ecto and transmembrane domains were fused to PD-1 tail and a C-terminal mGFP, denoted as BTLA^ECTO-TMD^-PD-1^CYTO^-mGFP (**Fig 3D**). This chimeric receptor was expressed at a similar level as BTLA (WT)-mGFP in their respective host Jurkat cells (**Fig 3E**), and contained a PD-1 tail that can be triggered by HVEM. Unlike BTLA(WT)-mGFP, which recruited both SHP1 and SHP2, this chimeric recruited only SHP2 (**Fig 3F**, GFP IP, SHP2 IB), supporting the model that BTLA but not PD-1 recruits SHP-1. Interestingly, the chimera, which harbors a PD-1 tail, recruited substantially more SHP2 than did BTLA(WT), confirming that SHP2 prefers PD-1 over BTLA, as shown in **Fig 2D**. Notably, in both experiments involving the chimeras, dephosphorylation of CD3ζ only occurred when the receptor contained an intact BTLA tail (**Fig 3C** and **F**, WCL, CD3ζ-pY142 IB). This result further supports the model that BTLA, but not PD-1, potently inhibits TCR phosphorylation via its association with SHP1.

### SHP1 dephosphorylates both TCR and CD28 while SHP2 dephosphorylates CD28

We next examined how genetic ablation of SHP1 and/or SHP2 from T cells affects the functions of PD-1 and BTLA. Using CRISPR/Cas9 technology, we generated SHP1 knockout (KO), SHP2 KO and SHP1/SHP2 double KO (DKO) Jurkat cells, and introduced either PD-1 (WT)-mGFP or BTLA (WT)-mGFP in these cells via lentiviral transduction (**Fig S3A** and **B**). In the SHP1 KO background (**Fig 4A**), both BTLA and PD-1 still inhibited CD28 phosphorylation, but the BTLA effect on CD3ζ phosphorylation was largely abolished (**Fig 4B**), even though SHP2 was recruited by BTLA (**Fig S4A**). This result indicates that SHP2 can substitute SHP1, to associate with BTLA, for CD28 dephosphorylation. However, SHP2, even coupled with BTLA, had limited effect on CD3ζ dephosphorylation. Hence, it is the ability of BTLA to recruit SHP1, rather than SHP2, that allows BTLA to inhibit TCR phosphorylation. Further, the IL-2 secretion qualitatively agrees with the phosphorylation readouts – in the SHP1 KO background, both PD-1 and BTLA were still able to inhibit IL-2 secretion (**Fig 4C**), perhaps due to their abilities to repress CD28 phosphorylation.

**Figure 4.**
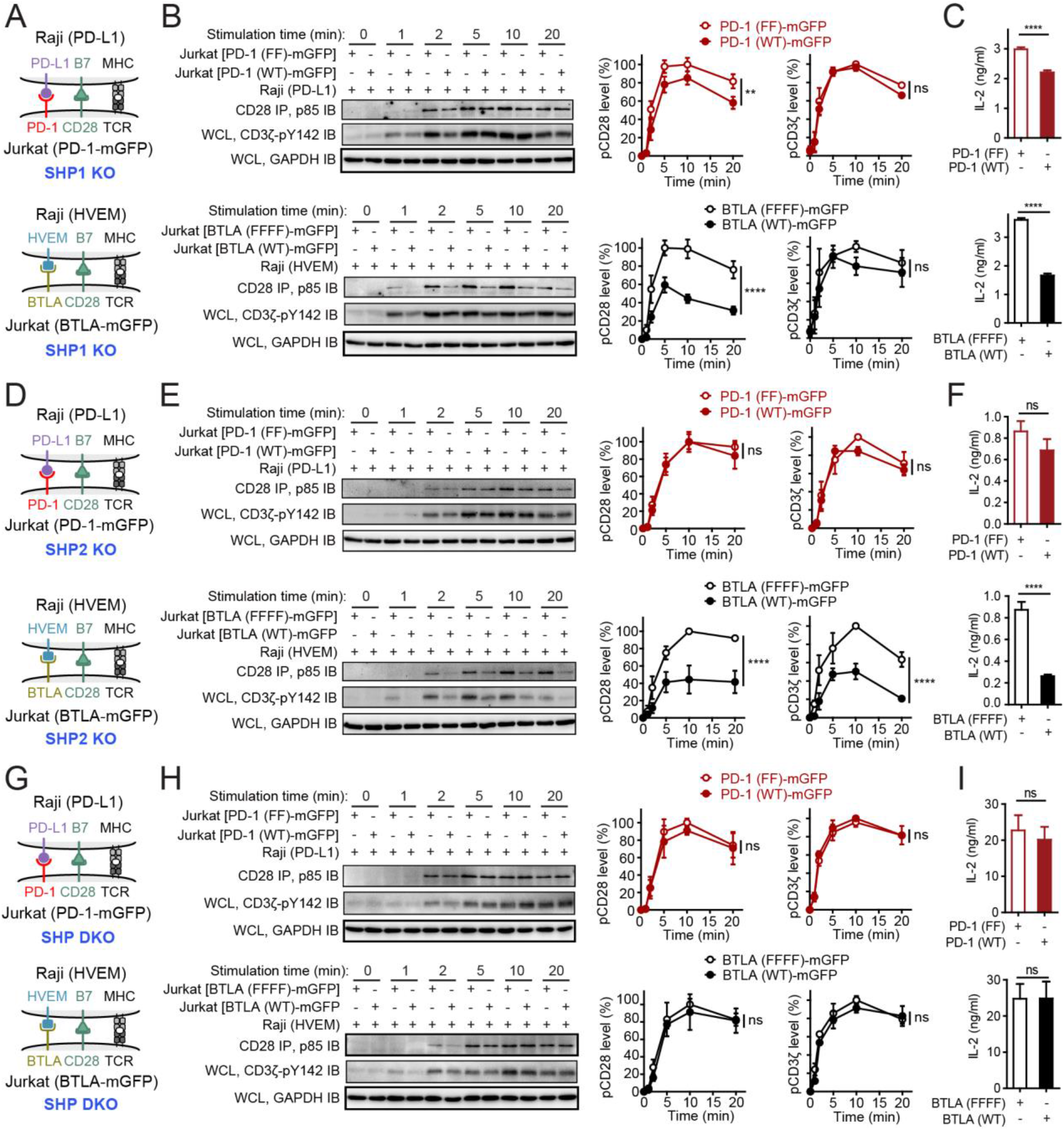
Effects of SHP1 and/or SHP2 KO on PD-1 and BTLA signaling. **(A)** Cartoons depicting a Raji-Jurkat co-culture assay to probe PD-1 or BTLA signaling in a SHP1 KO background. PD-1 (WT)-mGFP or PD-1 (FF)-mGFP transduced SHP1 KO Jurkat T cells were stimulated with PD-L1 or HVEM positive Raji B cells preloaded with SEE antigen. **(B)** Shown on the left are representative immunoblots showing the phosphorylations of CD3ζ (anti-pY142)and CD28 (co-IP’ed p85) for the indicated co-cultures with the cells lysed at the indicated time points after Raji-Jurkat contact (see **Methods**). Shown on the right are quantification graphs of CD28 and CD3ζ phosphorylations, incorporating results from three independent experiments. Data were normalized to the highest phosphorylation under PD-1 FF or BTLA FFFF condition, respectively. **(C)** Bar graphs summarizing the levels of secreted IL-2 of the indicated Raji-Jurkat co-cultures (see **Methods**). (**D-F**) Same as A-C except using SHP2 KO Jurkat cells. (**G-I**) Same as A-C except using SHP1/SHP2 double KO Jurkat cells. Data in this figure are presented as means ± SD from three independent measurements. ****P < 0.0001; ns, not significant; Two-way ANOVA (analysis of variance).

At the SHP2 KO background (**Fig 4D**), the PD-1 effect on CD28 was abolished, but the BTLA inhibitory effects on both CD3ζ and CD28 remained intact (**Fig 4E**). This result suggests that SHP2 is required for PD-1 to inhibit CD28, and SHP1 cannot substitute SHP2 to support PD-1 function. Indeed, no SHP1 recruitment to PD-1 was detected in the SHP2 KO background (**Fig S4B**). By contrast, strong BTLA:SHP1 co-IP was detected at this SHP2 KO background (**Fig S4B**), thereby allowing BTLA to execute its suppressive function. Hence, the lack of PD-1:SHP1 association at the WT Jurkat background (**Fig 2B** and **C**, **3C** and **F**) was likely due to an intrinsic weak propensity of SHP1 to bind PD-1, rather than a competition from SHP2. In agreement with the phosphorylation effects, SHP2 KO abolished the effect of PD-1, but not BTLA, on IL-2 production (**Fig 4F**).

At the DKO background (**Fig 4G**), both BTLA and PD-1 were phosphorylated upon Raji-Jurkat conjugation (**Fig S4C**), but neither inhibited CD3ζ or CD28 phosphorylation (**Fig 4H**), except perhaps a weak effect at two minutes, indicating that SHP1 and/or SHP2 are the major effectors for both BTLA and PD-1.

Taken together, results from the SHP1/SHP2 KO experiments demonstrate that BTLA:SHP1 complex can dephosphorylate both TCR and CD28, whereas BTLA:SHP2 or PD-1:SHP2 complex only dephosphorylate CD28 (**Fig 5**). We note that a recent study suggested that SHP2 is dispensable for PD-1 to cause T cell exhaustion (Rota et al., 2018), indicating that PD-1 might have other effectors. However, here we find that SHP2 KO completely abolishes the ability of PD-1 to inhibit CD28 phosphorylation and IL-2 secretion, suggesting that SHP2 is the major effector for PD-1 in T cells. Notably, SHP2 KO dramatically decreased the IL-2 secretion even for Jurkat expressing the signaling deficient mutants of PD-1 or BTLA (**Fig 4F** versus **1C** **and** **1F**). This result indicates that SHP2 is important in maintaining the function of T cells, independent of PD-1 or BTLA signaling. Indeed, previous studies have implicated SHP2 in activating Src family kinases (Zhang et al., 2004), and promoting T cell activation and development (Nguyen et al., 2006). Therefore, we speculate that SHP2 KO would impair T cell function, regardless the expression of PD-1, to produce an “exhaustion” like phenotype.

**Figure 5.**
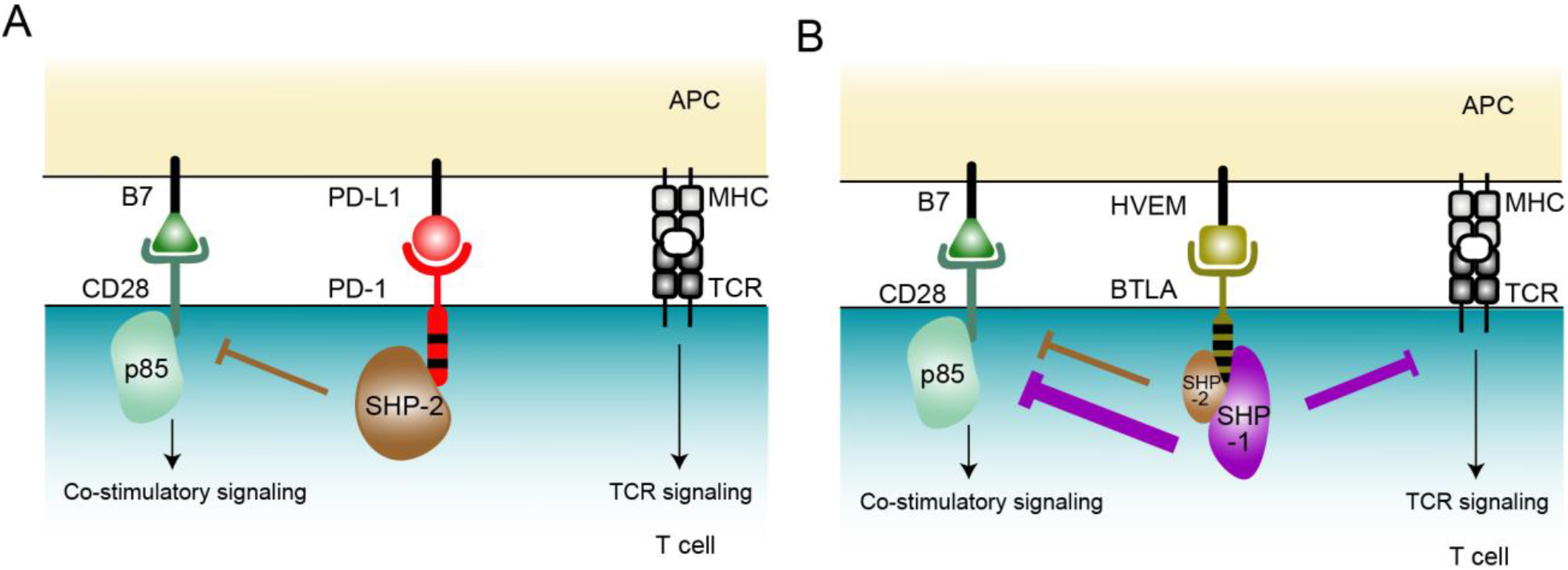
Proposed model for PD-1 and BTLA signaling. **(A)** PD-1 recruits SHP2, but not SHP1 to dephosphorylate CD28 (panel A). **(B)** BTLA recruits SHP1, and to a lesser extent, SHP2. The BTLA:SHP1 complex potently dephosphorylates both TCR and CD28, where the BTLA:SHP2 complex dephosphorylates CD28, with limited impact on TCR phosphorylation.

The unique ability of BTLA:SHP1 complex to dephosphorylate TCR suggests that SHP1 has an intrinsically higher phosphatase activity than SHP2. Consistent with this idea, a previous biochemical analysis showed that SHP1 displays 2-3 folds higher catalytic activity than does SHP2 towards a number of substrates (Ren et al., 2011).

In conclusion, we uncovered an unexpected disparity in the biochemical specificities of two structurally related immune checkpoints and two tyrosine phosphatase paralogs. While PD-1 couples SHP2 to preferentially dephosphorylate CD28 co-stimulatory receptor, BTLA employs SHP1, and to a less extent, SHP2 to dephosphorylate both TCR and CD28. This finding suggests that PD-1 and BTLA have evolved to divide work tasks in repressing the T cell response. Our finding that SHP1 but not SHP2 targets ITAM of TCR have implications for the functions of these two phosphatases, as well as receptors that recruit these phosphatases. It is likely that SHP1-recruiting receptors are responsible for dephosphorylating ITAM-containing receptors, while SHP2-recruiting receptors primarily dephosphorylate non-ITAM receptors. Future studies are needed to test the generality of this model.

## MATERIALS AND METHODS

### Reagents

RPMI 1640 (#10-041-CM) was purchased from corning. DMEM (#25-501) was purchased from Genesee Scientific. 100× Penicillin-Streptomycin solution (#SV30010) was obtained from GE Healthcare. Fetal bovine serum was obtained from Omega Scientific (#FB-02). Lipofectamine Transfection Reagent (#18324010) was purchased from Thermo Fisher Scientific. D-Desthiobiotin (#D1411) and PY20 monoclonal anti-phospho-tyrosine antibody (#P4110) were purchased from Sigma-Aldrich. SEE (#ET404) super antigen was purchased from Toxin Technology. CD28 monoclonal antibody (#16-0289-85), GFP polyclonal antibody (#A6455), PD-1 PE (#12-9969-42) and Dynabeads Protein G (#10004) were purchased from Thermo Fisher Scientific. PI3 Kinase p85 polyclonal antibody (#4292S) was obtained from Cell Signaling Technology. Anti-CD3ζ (pY142) monoclonal antibody (#558489) was obtained from BD Biosciences. GFP-Trap (#gta-20) was purchased from Chromotek. SHP1 monoclonal antibody (#sc-7289), SHP2 monoclonal antibody (#sc-7384), SAP monoclonal antibody (#sc-393948) and GRB2 monoclonal antibody (#sc-8034) were obtained from Santa Cruz Biotechnology. GAPDH polyclonal antibody (#10494-1-AP) was purchased from ProteinTECH. Antibodies of BTLA PE (#344505), CD28 APC (#302911), HVEM APC (#318807), PD-L1 APC (#393609), and ZAP-70 monoclonal antibody (#313402) were purchased from BioLegend.

### Cell cultures

Jurkat T cells were obtained from Dr. Arthur Weiss (University of California San Francisco). HEK293T cells and Raji B cells were obtained from Dr. Ronald Vale (University of California San Francisco). HEK293T cells were maintained in DMEM medium (DMEM supplemented with 10% fetal bovine serum, 100 U/mL of Penicillin, and 100 μg/mL of Streptomycin) at 37°C / 5% CO_2_. Jurkat and Raji cells were maintained in RPMI medium (RPMI 1640 supplemented with 10% fetal bovine serum, 100 U/mL of Penicillin, and 100 μg/mL of Streptomycin) at 37°C / 5% CO_2_.

### Recombinant proteins

Human protein tyrosine kinase Lck (with a G2A mutation to abolish myristoylation) was expressed and purified with an N-terminal His10 tag as described (Hui and Vale, 2014). The cytosolic tails of PD-1 WT (aa 194-288), PD-1 FF mutant (aa 194-288, Y223, 248F), BTLA WT (aa 190-289), and BTLA FFFF mutant (aa 190-289, Y226, 243, 257, 282F) were all expressed with an N-terminal Glutathione S-transferase (GST)-tag in *Escherichia coli* using the pGEX-6p2 vector. Human full-length protein tyrosine phosphatase SHP1 and SHP2 were expressed with an N-terminal GST tag followed by a PreScission recognition sequence (LEVLFQGP), and a SNAP-Tag in the Bac-to-Bac baculovirus expression system. All GST fusion proteins were purified using Glutathione Sepharose 4B. All poly-histidine tagged proteins were purified using the Ni-NTA agarose (GE Healthcare), and eluted with imidazole. All affinity purified proteins were subjected to gel filtration chromatography using HEPES buffered saline (50 mM HEPES-NaOH, pH 7.5, 150 mM NaCl, 10% glycerol, 1 mM TCEP). The monomer fractions were pooled, aliquoted, snap frozen and stored at −80 °C.

### Cell lines generation

To generate SHP1, or SHP2 KO Jurkat cells, two PX330-GFP vectors coding SHP1, or SHP2 specific sgRNA were electroporated into Jurkat cells with Bio-Rad Gene Pulser Xcell by exponential protocol (250 V, 1000 μF, ∞ Ω, 4 mm cuvette). To generate SHP1 and SHP2 DKO Jurkat cells, two PX330-GFP vectors coding SHP1 sgRNA were electroporated into SHP2 KO Jurkat cells by the same way. After electroporation, cells were recovered in culture medium for two days at 37°C / 5% CO_2_. Single GFP positive cells were then sorted to 96-well plate with FACS and maintained in culture media for three weeks, after which the corresponding KO cell clones were validated by western blot with anti-SHP1 and anti-SHP2 antibodies.

Each gene of interest was introduced into Jurkat and Raji cells via lentiviral transduction, as described previously (Zhao et al., 2018). WT Raji were lentivirally transduced with PD-L1– mCherry or HVEM–mRuby2 to generate Raji (PD-L1) and Raji (HVEM). WT, SHP1 KO, SHP2 KO, SHP DKO Jurkat cells with PD-1 WT, PD-1 FF, BTLA WT, or BTLA FFFF were generated by transducing PD-1 WT–mGFP, PD-1 FF–mGFP, BTLA WT–mGFP, or BTLA FFFF–mGFP into Jurkat cells at WT, SHP1 KO, SHP2 KO, or SHP1/2 DKO background (Hui et al., 2017). Jurkat (PD-1^ECTO-TMD^-BTLA^CYTO^) was generated by lentivirally transducing Jurkat WT with PD-1^ECTO-TMD^-BTLA^CYTO^–mGFP. Jurkat (BTLA^ECTO-TMD^-PD-1^CYTO^) was generated by transducing Jurkat WT with BTLA^ECTO-TMD^-PD-1^CYTO^–mGFP via lentivirus under the control of the dSV40 promoter.

### Jurkat-Raji conjugation assay

For cell conjugation assay, Jurkat cells were starved in serum free RPMI medium at 37°C for three hours before conjugation to reduce the phosphorylation background. Raji B cells were pre-incubated with 30 ng/mL SEE (Toxin Technology) in RPMI medium for 30 minutes at 37°C. Following antigen loading, 2 million antigen-loaded Raji B cells and 2 million Jurkat T cells were precooled on ice and mixed in a 96-well plate. The plate was then centrifuged at 290× g for one minute at 4 °C to initiate cell–cell contact, and immediately transferred to a 37°C water bath. The reactions were terminated with NP-40 lysis buffer (50 mM HEPES, pH 7.4, 150 mM NaCl, 1% NP-40, 1 mM EDTA, 5% glycerol, 1 mM PMSF, 10 mM Na_3_VO_4_, 10 mM NaF) at indicated time points. CD28 was co-immunoprecipitated from the lysate using anti-CD28 antibody (Thermo Fisher Scientific, #16-0289-85). BTLA–mGFP and PD-1–mGFP were co-immunoprecipitated from the lysate using GFP-Trap (Chromotek, #gta-20). Equal fractions of the immunoprecipitates were subjected to SDS-PAGE and indicated antibodies.

### IL-2 ELISA assay

Jurkat cells were starved in serum free RPMI medium at 37°C for three hours before conjugation to reduce the phosphorylation background. Raji B cells were pre-incubated with 30 ng/mL SEE superantigen (Toxin Technology) in RPMI medium for 30 minutes at 37°C. Following antigen loading, 0.1 million antigen-loaded Raji B cells and 0.2 million Jurkat T cells were mixed in a 96-well U-bottom plate in triplicate wells. The plate was then centrifuged at 290× g for one minute to initiate cell–cell contact, and immediately transferred to 37°C / 5% CO_2_ incubator. Supernatants were collected at 6 hours. Human IL-2 was quantified by ELISA (BioLegend, # 431804).

### Flow cytometry

Flow cytometry was carried out using either LSRFortessa cell analyzer (BD Biosciences) or FACSAria Fusion flow cytometer (BD Biosciences) and data analyzed using FlowJo software (FlowJo, LLC). For **Fig S1**, Raji cells were stained with APC-labeled anti-HVEM (or isotype), or APC-labeled anti-PD-L1 (or isotype) to detect HVEM or PD-L1 expression. For **Fig 1B, 1E**, and **S3B**, Jurkat cells were stained with PE-labeled anti-BTLA (or isotype), PE-labeled anti-PD-1(or isotype), or APC-labeled anti-CD28(or isotype) to detect BTLA, PD-1, or CD28 expression, respectively.

### GST pull down assay

The GST-tagged baits (10 μg) were pre-phosphorylated by 0.2 μg His-Lck (G2A) at room temperature for one hour, then mixed with the Raji stimulated Jurkat cell lysates for 3 h at 4 °C, then added 100 μL pre-equilibrated Glutathione agarose resin (GoldBio, G-250) and rotated for one hour at 4 °C. Beads were wash with 1 mL wash buffer (20 mM HEPES, pH 7.4, 50 mM NaCl, 0.1% NP-40, 1 mM EDTA, 10 mM Na_3_VO_4_, 10 mM NaF) for 4 times, proteins were eluted off the beads by boiling with SDS loading buffer.

### Liquid chromatography-tandem mass spectrometry (LC-MS/MS)

The samples were digested in gel and analyzed on LC-MS as described (Peled et al., 2018). The MS/MS spectra were searched against the UniProt Human reference proteome database (downloaded June 2^nd^, 2018), with WT and signaling-deficient mutant GST-BTLA (aa 190-289), GST–PD-1(aa 190-288) sequences inserted into the database, using SEQUEST within Proteome Discoverer. Database queries to sort for SH2-containing proteins were conducted as described in (Peled et al., 2018).

### Quantification and Statistical Analysis

Data were shown as mean ± SEM or mean ± SD, and number of replicates were indicated in figure legends. Curve fitting and normalization were performed in GraphPad Prism 5. Statistical significance was evaluated by two-way ANOVA (*, p < 0.05; **, p < 0.01; ***, p < 0.001; ****, p < 0.0001) in GraphPad Prism 5. Data with p < 0.05 are considered statistically significant.

## AKNOWLEDGEMENTS

We thank X. Chen and T. Masubuchi for critically reading the manuscript. This work was supported by NIH grant R37 CA239072 to E. Hui, NSFC grants 31770952 & 31570911 to G. Fu, and NSFC grant 31670919 to H. Wang. Y. Zhao is a Cancer Research Institute Irvington Postdoctoral Fellow. H. Wang is funded by the 1,000-Youth Elite Program of China. E. Hui is a Searle Scholar and a Pew Biomedical Scholar.

---

**Figure S1.**
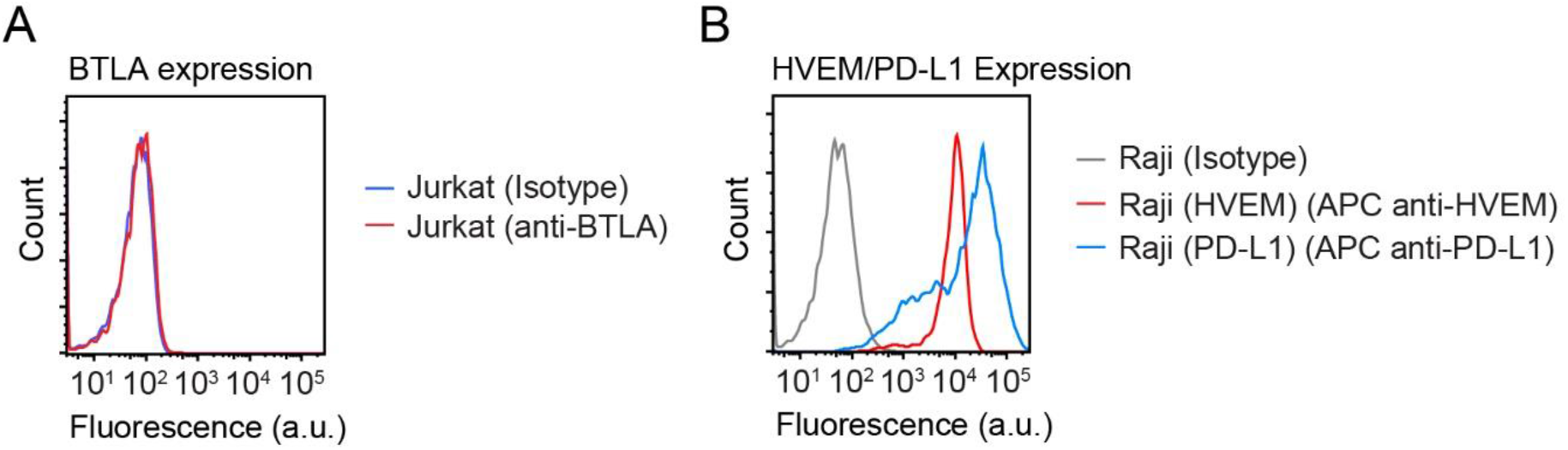
Jurkat T cells do not express BTLA. **(A)** Shown in red is a flow cytometry histogram of BTLA surface expression on parental Jurkat cells. Shown in blue is the histogram for the isotype control staining. **(B)** Shown in red is a flow cytometry histogram of HVEM surface expression on Raji (HVEM) cells. Shown in blue is a flow cytometry histogram of PD-L1 surface expression on Raji (PD-L1) cells. Shown in grey is a histogram for the isotype control staining. a. u. denotes arbitrary units.

**Figure S2.**
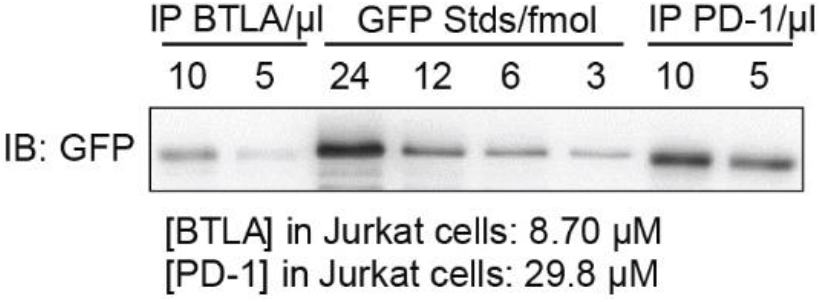
Quantification of BTLA-mGFP and PD-1-mGFP in Jurkat (BTLA-mGFP) and Jurkat (PD-1-mGFP) cells respectively. 10 million of either Jurkat (BTLA-mGFP) or Jurkat (PD-1-mGFP) cells were lysed with PBS containing 1% NP40 with protease inhibitors on ice. BTLA-mGFP and PD-1-mGFP were then immunoprecipitated using GFP-trap, and eluted by 50 μl of SDS sample buffer. 10 μl and 5 μl of the eluates were loaded to SDS-PAGE together with indicated amounts of purified GFP standard (std), and transferred to nitrocellulose membranes, and subjected to immunoblot (IB) with an anti-GFP antibody. BTLA-mGFP and PD-1-mGFP were then quantified using a standard curve constructed based on the GFP standards. Finally, the BTLA and PD-1 concentrations in Jurkat cells were calculated by dividing their moles by the average volume of a Jurkat cell, assuming an average diameter of 12 μm for Jurkat cells.

**Figure S3.**
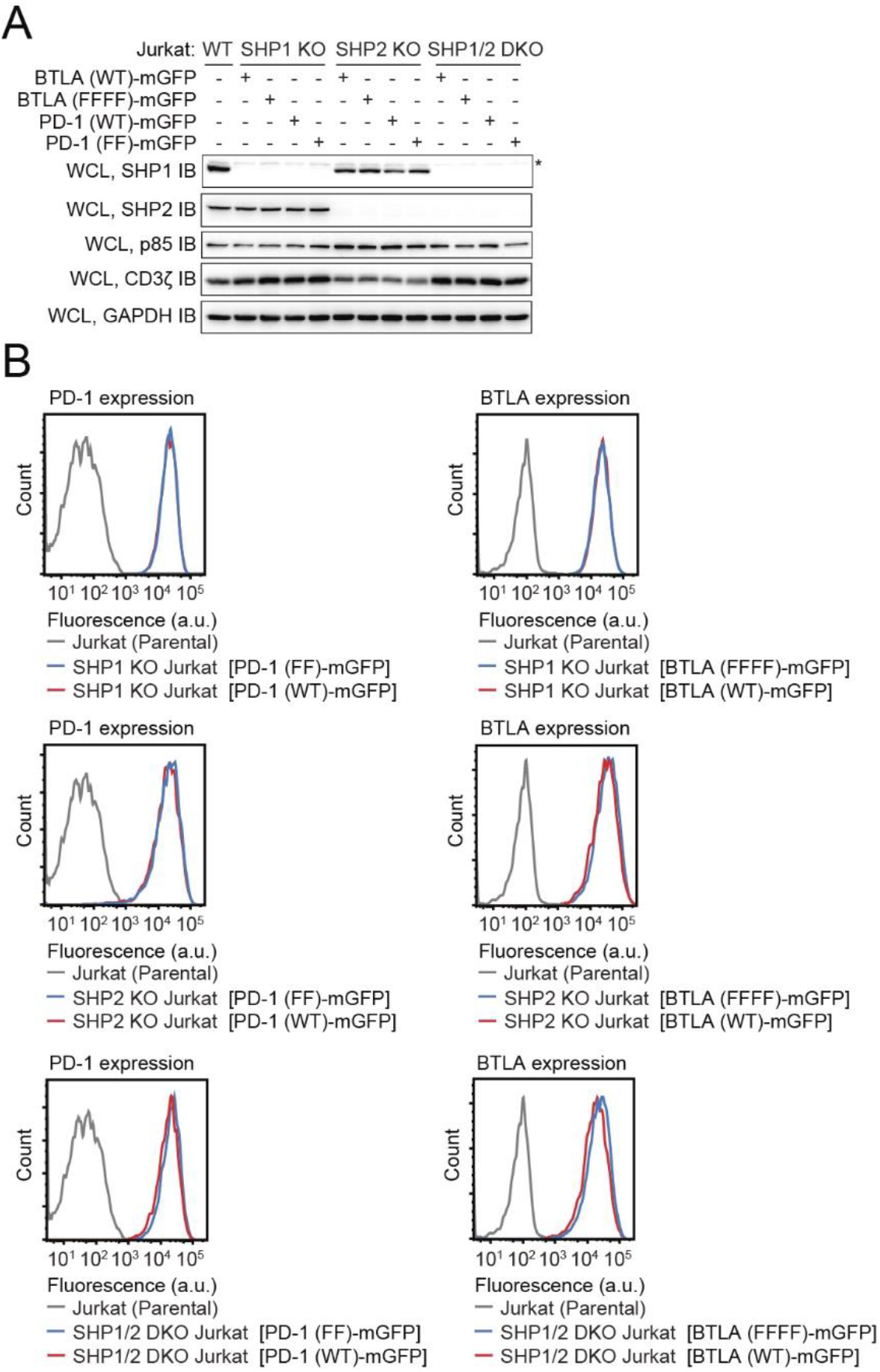
Expression levels of SHP1, SHP2, BTLA, and PD-1 on SHP1 KO, SHP2 KO, or DKO Jurkat cells. **(A)** Immunoblots showing the levels of SHP1, SHP2, p85 and CD3ζ in WT Jurkat cells or BTLA (WT), BTLA (FFFF), PD-1 (WT) or PD-1 (FF)-transduced SHP1 KO, SHP2 KO or SHP DKO Jurkat cells. GAPDH blots indicate the input of each sample. Non-specific bands were labeled by asterisks. **(B)** Flow cytometry histograms showing BTLA or PD-1 surface expression in parental Jurkat cells and BTLA or PD-1 transduced SHP1 KO, SHP2 KO, SHP1/2 DKO Jurkat cells. a. u. denotes arbitrary units.

**Figure S4.**
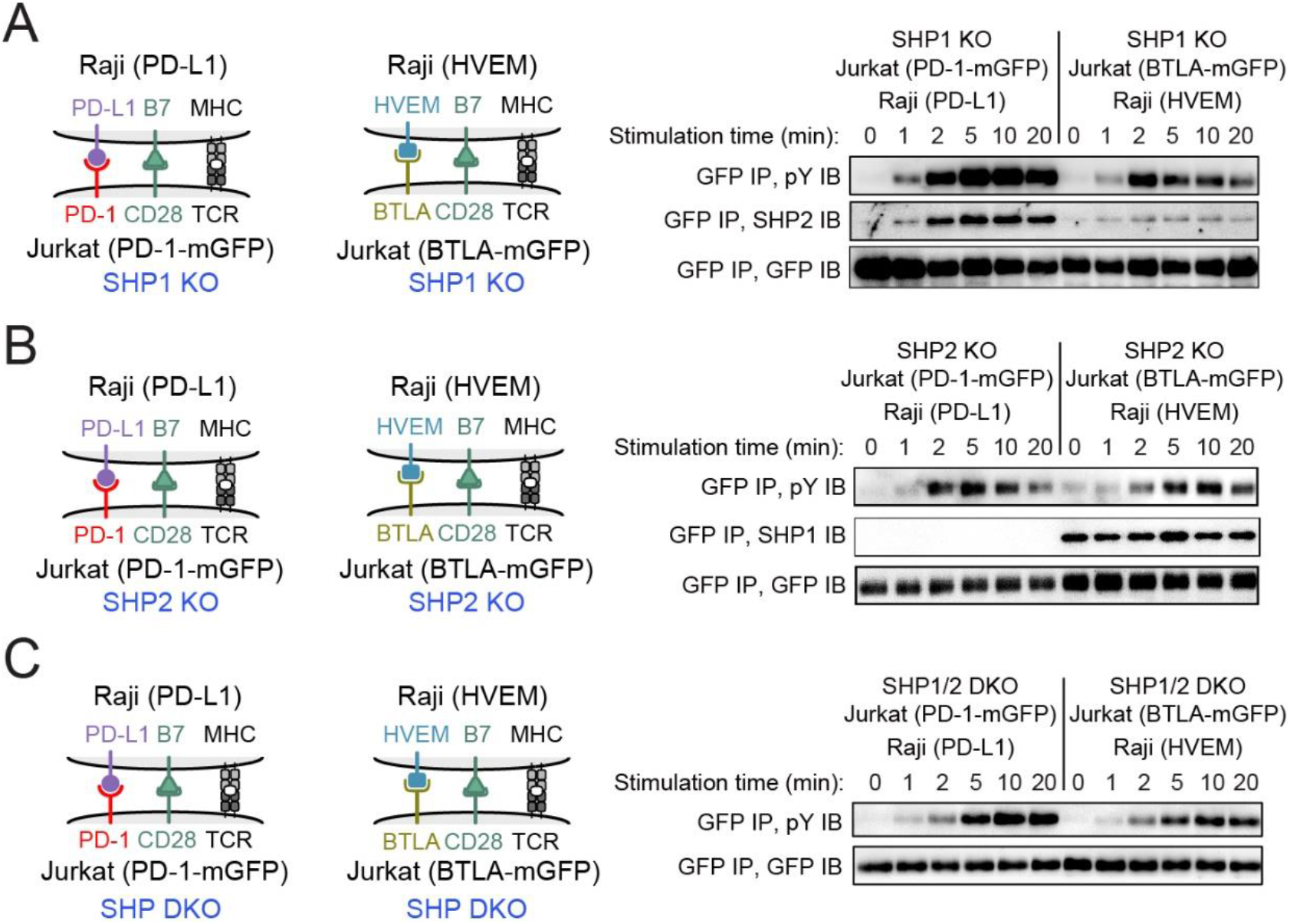
Recruitment of SHP1/2 by BTLA or PD-1 in SHP2 KO or SHP1 KO Jurkat cells. **(A-C)** Immunoblots showing PD-1 or BTLA co-precipitated SHP1 or SHP2 in the lysates of the indicated co-cultures, with the stimulation time (duration of cell-cell contact before lysis) indicated (see **Methods**). pY, immunoblot for phospho-tyrosine on PD-1 or BTLA by pY20 monoclonal antibody. GFP was immunoblotted to show the input of PD-1-mGFP or BTLA-mGFP.

